# BiocPy: A Framework for Cross-Platform Genomic Data and Methods Interoperability

**DOI:** 10.64898/2026.11.14.766721

**Authors:** Jayaram Kancherla, Aaron T.L. Lun

**Affiliations:** Genentech Inc., South San Francisco, CA, USA; Independent Scientist, Sydney, Australia

## Abstract

Bioconductor provides standardized data structures and statistical methods for high-throughput genomic analysis in R, whereas Python has become a primary environment for machine learning, image analysis, and general-purpose data science. Combining tools across both ecosystems often requires ad-hoc file conversions or embedded language runtimes that can lose metadata, introduce maintenance overhead, or lead to subtle differences in downstream results. Here, we present BiocPy, a collection of Python packages that support cross-language genomic workflows through three complementary layers: (1) native Python implementations of Bioconductor’s core data structures; (2) lossless data exchange via direct reading and writing of serialized R `(.rds)` objects and the language-neutral `ArtifactDB` format for persistent, multi-stage pipelines; and (3) a cross-platform method development paradigm built on shared C++ libraries wrapped across R, Python, and WebAssembly to support identical statistical analyses across environments. All packages in the BiocPy ecosystem are published to the Python Package Index (PyPI) with source code hosted at https://github.com/BiocPy.

## 1 Introduction

Bioinformatics sits at the intersection of molecular biology, statistics, and computer science. Bioconductor[1, 2], built upon the R programming language, was designed with a focus on statistical analysis and visualization of genomic data. Over the past two decades, it has grown into a repository of over 2,000 packages organized around shared data structures and design principles. Classes such as `GenomicRanges`[3] and `SummarizedExperiment`[4] serve as a common currency for data exchange, allowing developers to build and compose methods without defining bespoke representations for each package. Bioconductor remains a standard environment for statistical testing and established genomic workflows.

Over the same period, Python has become the primary language for general-purpose data science and machine learning (ML). Supported by libraries such as NumPy[5], PyTorch[6], and TensorFlow[7], Python offers broad integration with modern computing infrastructure, including distributed computing and GPU acceleration. In genomics, Python adoption has grown substantially with single-cell analysis frameworks such as scverse[8] and GPU-accelerated toolkits such as rapids-singlecell[9].

In practice, genomic analyses frequently span both ecosystems: an analyst may rely on Bioconductor packages for experimental design, normalization, or differential expression, and then move to Python for deep learning or large-scale integration. Bridging these steps typically relies on exporting intermediate flat files, which strips nested metadata and requires manual reassembly, or reimplementing statistical routines in Python, which can introduce subtle algorithmic discrepancies.

BiocPy addresses this gap by implementing Bioconductor’s core class definitions directly in Python. Rather than embedding an R session inside Python or proposing a new data standard, BiocPy provides native Python classes such as `GenomicRanges` and `SummarizedExperiment` that mirror the structure and behavior of their R counterparts. Keeping the conceptual data model consistent reduces friction when moving data or translating workflows between languages, while still allowing direct conversion to standard Python structures when needed.

The BiocPy framework is targeted at developers who require the reliability of Bioconductor’s data structures within a Python environment. Unlike rpy2[10], which embeds an R runtime and can complicate deployments, BiocPy is lightweight and native to Python. Similarly, while `AnnData`[11] is well-suited for single-cell expression matrices, it lacks the broader generality of `SummarizedExperiment` and `MultiAssayExperiment` and does not natively support genomic interval coordinates (such as `GenomicRanges`) or range-based operations. BiocPy supports this broader spectrum of genomic data types while providing converters to and from `pandas, AnnData`, and `MuData`.

## 2 Core Data Structures

BiocPy adapts Bioconductor’s S4 design principles to idiomatic Python rather than simply translating the R class definitions. The goal is not to replace existing Python scientific libraries, but to provide containers that maintain structural and semantic equivalence with Bioconductor objects. Although BiocPy classes provide methods to convert to and from popular Python representations such as pandas `DataFrames`[12] or `AnnData`, those representations do not always capture nested metadata, genomic range coordinates, or sample-level mappings without loss of information. BiocPy containers preserve these members so that objects can move between R and Python without losing structural fidelity.

A central design difference between R and Python is object mutability. R uses copy-on-write semantics, which prevents modifications to an object inside a function from unexpectedly mutating references held elsewhere. Python passes object references by default; in-place modifications to a shared metadata table or coordinate vector can silently corrupt parent containers or bypass dimensional validation. To prevent unintended side effects in multi-step pipelines, BiocPy encourages a functional programming style: setter and transformation methods (e.g., `set_column()`, `set_row_names()`, `set_assay()`) default to `in_place=False`, returning a new shallow copy that shares unmodified underlying arrays while updating only the affected member. For interactive convenience, standard Python property assignments and item setters are also supported as aliases for in-place updates (`in_place=True`), which still trigger container validation checks.

At its core, the `biocutils` package provides S4-like generic functions using Python’s `singledispatch` mechanism, such as `combine_rows()`, `combine_columns()`, `subset()` etc., along with base 1D data structures such as `Factor`, `Names`, and typed list classes (`StringList`, `IntegerList`, `FloatList`, `BooleanList`). Core numerical representations build directly on NumPy and SciPy[13], keeping the dependency footprint minimal. Table 1 summarizes the primary packages in the BiocPy ecosystem and their Bioconductor counterparts.

**Table 1:** Overview of core packages in the BiocPy ecosystem, their Bioconductor equivalents, and their primary functionality.

| **BiocPy Package** | **Bioconductor Equivalent** | **Primary Role** |
|---|---|---|
| biocutils | BiocGenerics, S4Vectors | Generic functions (combine_rows, subset), Factor, Names, and typed lists |
| biocframe | S4Vectors::DFrame | Metadata table supporting nested and arbitrary column objects |
| iranges, genomicranges | IRanges, GenomicRanges | 1D and genomic interval containers, set operations, and overlap search |
| compressed-lists | IRanges::CompressedList | Contiguous partitioned storage for grouped vectors and ranges |
| summarizedexperiment | SummarizedExperiment | Matrix-like assay containers (SE and RSE) linked to feature and sample metadata |
| singlecellexperiment | SingleCellExperiment | Single-cell container adding reduced_dims and alt_exps |
| multiassayexperiment | MultiAssayExperiment | Multi-omics cohort container coordinating experiments through a sample map |
| delayedarray | DelayedArray, HDF5Array | Lazy array evaluation over on-disk backends (HDF5, TileDB) |
| rds2py, dolomite-* | readRDS / saveRDS, alabaster.* | Direct R .rds reading/writing and language-agnostic ArtifactDB serialization |
| scranpy, singler, scrnaseq, celldex | scrapper, SingleR, scRNAseq, celldex | Single-cell analysis, reference annotation, and curated datasets |
| txdb, orgdb, ensembldb, experimenthub | GenomicFeatures, AnnotationDbi, ensembldb, ExperimentHub | Transcript, gene, and organism annotation queries and hub resource access |

### 2.1 BiocFrame: A Flexible DataFrame

The `BiocFrame`[14] class implements the semantics of Bioconductor’s `DFrame` (`DataFrame`) class from `S4Vectors`[15] and serves as the primary metadata container across the BiocPy ecosystem. While pandas `DataFrames` are standard in Python data analysis, they enforce columnar type homogeneity backed by NumPy or Arrow arrays. In genomic workflows, this can lead to unwanted type coercion (such as casting integers to floats when missing values are introduced) and makes it difficult to store complex domain objects as columns.

In contrast, `BiocFrame` makes minimal assumptions about its columns: any object that defines a length (`__len__`) equal to the number of rows and supports slicing (`_getitem_`) can be stored as a column. This protocol-based design preserves original data types and allows arbitrarily nested or specialized columns such as another `BiocFrame`, a 2D NumPy array, a `GenomicRanges` object, or a compressed list of intervals, to be sliced and combined alongside standard 1D vectors. `BiocFrame` also supports Bioconductor’s metadata members, including `column_data` (a `BiocFrame` describing the columns) and `metadata` (an arbitrary dictionary describing the table). When users need general tabular operations or plotting utilities, `BiocFrame` provides direct conversion methods (`to_pandas()`, `from_pandas()`, `to_polars()`, `from_polars()`) to interoperate with pandas and Polars[16], though converting to flat tabular formats may omit nested metadata or specialized column types.

### 2.2 GenomicRanges and Interval Representations

Interval arithmetic on genomic coordinates is a core requirement of many sequencing workflows. In Bioconductor, genomic intervals are represented using the `IRanges` and `GenomicRanges`[3] packages. In Python, interval operations have typically been performed by wrapping command-line utilities such as bedtools[17] or using tabular libraries such as PyRanges[18]. BiocPy provides native, in-memory implementations of `IRanges` and `GenomicRanges`[19] that follow Bioconductor’s coordinate and strand semantics.

Unlike a generic table with chromosome, start, and end columns, a `GenomicRanges` object couples integer intervals (via an `IRanges` instance storing start positions and widths) with sequence names (`seqnames`), strand orientation (+, −, or *), feature-level metadata (`mcols`, stored as a `BiocFrame`), and reference genome metadata (`SeqInfo`, tracking chromosome lengths and circularity). This structure enforces coordinate validity and strand-aware arithmetic across operations including overlap detection (`find_overlaps()`, `count_overlaps()`, `subset_by_overlaps()`), proximity queries (`nearest()`, `precede()`, `follow()`), intra-range transformations (`shift()`, `resize()`, `flank()`, `promoters()`), and inter-range set operations (`reduce()`, `gaps()`, `disjoin()`). For example, calling `promoters()` on gene annotations automatically accounts for strand direction when computing upstream and downstream coordinates relative to transcription start sites.

To represent grouped genomic intervals such as exons grouped by transcript or peaks grouped by sample, BiocPy also implements `CompressedGenomicRangesList` (`GenomicRangesList`). Built on the `CompressedList` abstraction, all constituent ranges are stored in a single contiguous `GenomicRanges` object alongside an integer partitioning vector. This avoids the memory and iteration overhead of instantiating thousands of separate Python objects while allowing vectorized operations across groups.

To support interval queries in Python, `iranges` and `genomicranges` delegate overlap and proximity searches to a C++ implementation of the nested containment list data structure[20] (`nclist-cpp`), with built-in multi-threading support (`num_threads`). Table 2 compares the runtime of `find_overlaps()` and `nearest()` in BiocPy against R’s `GenomicRanges` (v1.64.0) across 10^5^ to 10^6^ randomly sampled genomic intervals distributed over 24 human chromosomes (evaluated on an Intel Core i7-8850H CPU at 2.60 GHz, averaged over 3 runs). Single-threaded BiocPy queries execute in under a second even for 10^6^ × 10^6^ interval comparisons, outperforming R’s GenomicRanges by 2× to 10×, with further speedups when multi-threading is enabled.

**Table 2:**
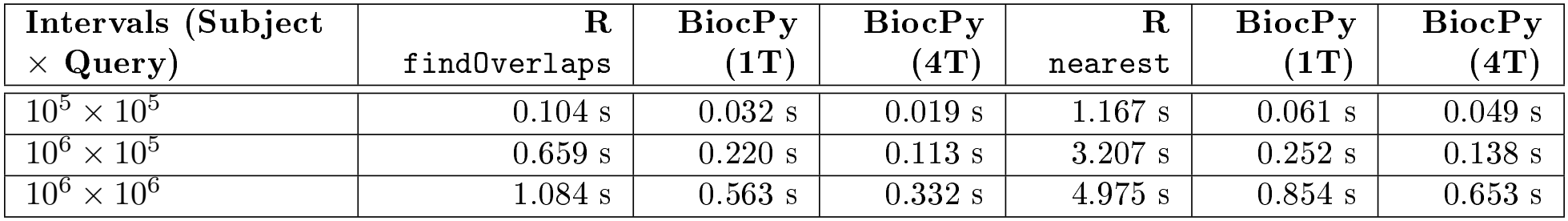
Wall-clock runtime (in seconds, mean of 3 runs) for genomic interval overlap (find_overlaps) and proximity (nearest) queries across 24 chromosomes, comparing R GenomicRanges against BiocPy GenomicRanges (1 and 4 threads).

### 2.3 Containers for Experimental Data

To organize high-throughput experimental assays alongside feature and sample annotations, BiocPy implements a hierarchy of experimental containers that mirror Bioconductor’s S4 classes (Figure 1). Every container validates dimensional consistency on construction and modification, ensuring that subsetting rows or columns remains synchronized across all assay matrices, feature annotations, and sample metadata.

**Figure 1:**
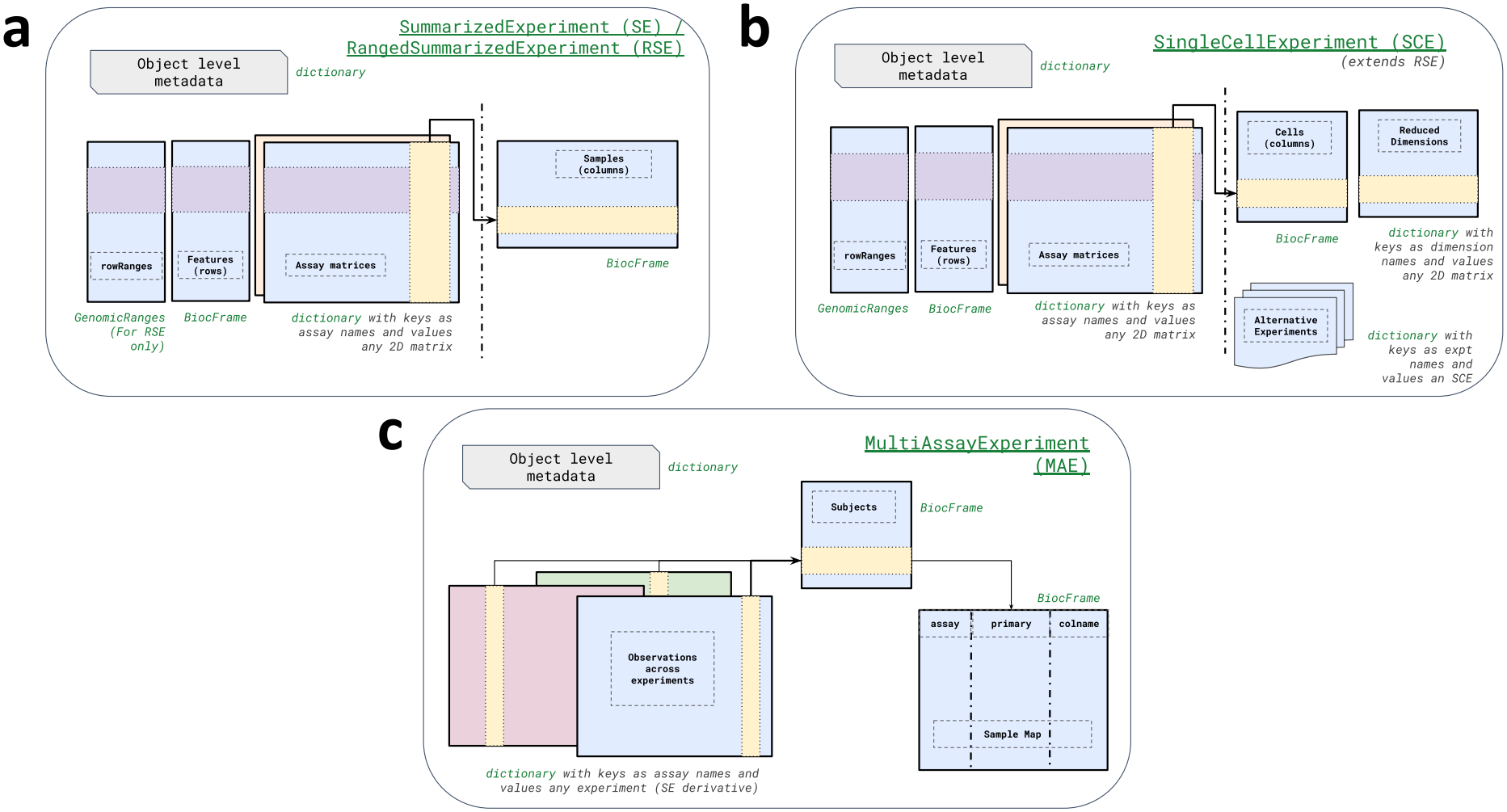
Class hierarchy and data members for experimental containers designed to mirror Bioconductor S4 semantics. a) `SummarizedExperiment` (SE) and `RangedSummarizedExperiment` (RSE) link assay matrices to row (genomic features) and column (sample) metadata. b) `SingleCellExperiment` (SCE) extends the RSE to include members for reduced dimensions (e.g., PCA, UMAP) and alternative experiments (e.g., spike-ins, ADTs). c) `MultiAssayExperiment` (MAE) coordinates multiple experiments across a cohort, utilizing a sample map to synchronize subsetting operations across diverse biological assays.

#### 2.3.1 SummarizedExperiment (SE) & RangedSummarizedExperiment (RSE)

`SummarizedExperiment`[21] serves as the base container for rectangular experimental data, where rows correspond to features (such as genes, transcripts, or genomic regions) and columns correspond to samples or cells. An SE instance stores one or more identically shaped 2D matrices in its `assays` dictionary (supporting dense NumPy arrays, SciPy sparse matrices, or `DelayedArray` objects), paired with `row_data` and `column_data` members represented as `BiocFrame` objects. `RangedSummarizedExperiment` extends SE by adding a `row_ranges` member (holding a `GenomicRanges` or `GenomicRangesList` object), enabling range-based queries (such as subsetting an expression matrix by genomic overlap) directly on the container. Note that this feature-by-sample orientation is the transpose of the observations-by-features layout used in `AnnData`. Both SE and RSE provide conversion methods to and from `AnnData`, however, this conversion is lossy because AnnData does not natively preserve `GenomicRanges` or `GenomicRangesList` coordinates (`row_ranges`), nested `BiocFrame` columns, or arbitrary container metadata required for lossless round-tripping with Bioconductor. In addition, `AnnData` does not accept arbitrary matrix representations, so converting an SE or RSE backed by a `DelayedArray` (described in the next section) forces those assays to be realized into memory.

#### 2.3.2 SingleCellExperiment (SCE)

Extending RSE, `SingleCellExperiment`[22, 23] adds members tailored for single-cell workflows. The `reduced_dims` member stores low-dimensional embeddings (such as PCA, t-SNE, or UMAP matrices), while the `alternative_experiments` (`alt_exps`) member stores auxiliary `SummarizedExperiment` objects that share the same cells (columns) but measure distinct features, such as antibody-derived tags (ADTs) in CITE-seq, CRISPR guide counts in Perturb-seq, or spike-in controls in historical scRNA-seq experiments. Subsetting cells in the parent SCE automatically subsets all reduced dimensions and alternative experiments. Because SCE derives from RSE, it likewise provides conversion methods to and from `AnnData` (mapping assays, cell and feature annotations, and `reduced_dims`), though the conversion remains lossy: `AnnData` cannot natively represent `row_ranges`, `alt_exps` with distinct feature spaces, or nested `BiocFrame` metadata needed for lossless Bioconductor round-tripping, and similarly realizes delayed assay matrices into memory.

#### 2.3.3 MultiAssayExperiment (MAE)

BiocPy provides `MultiAssayExperiment`[24, 25] for cohort studies where multiple experimental modalities (e.g., bulk RNA-seq, somatic variants, methylation, or proteomics) are measured across a shared set of biological subjects. Unlike single-cell multimodal containers where modalities are assumed to share identical columns, clinical and multi-omics cohorts often have missing assays, technical replicates, or multiple specimens per patient. MAE manages a dictionary of `experiments` (each an SE or SCE), patient-level `column_data`, and a `sample_map` (a `BiocFrame` linking each subject identifier to specific assay column names), supporting coordinated subsetting by feature, subject, or assay.

To interoperate with the broader Python single-cell ecosystem, MAE can be converted either to multiple `AnnData` objects (one per experiment) or to a unified `MuData`[26] container. However, both conversions are lossy because neither `AnnData` nor `MuData` natively encodes MAE’s `sample_map` for non-one-to-one, replicate, or missing sample-to-assay relationships. Retaining the native BiocPy classes therefore remains advantageous when workflows require range-based operations on `row_ranges`, coordinated cohort-level subsetting via `sample_map`, lazy delayed assay matrices, or lossless round-tripping with Bioconductor.

### 2.4 Delayed Arrays

To support datasets that exceed available RAM, BiocPy implements lazy array operations modeled on Bioconductor’s `DelayedArray`[27, 28] framework. Any experimental container in Section 2.3 can store `DelayedArray` objects directly in its `assays` member. Rather than executing array operations immediately, `DelayedArray` records subsetting, transposition, combining, and element-wise mathematical operations as delayed operations on a file, such as an on-disk HDF5[29] dataset or TileDB[30] array.

`DelayedArray` objects are conceptually similar to Dask arrays[31] and can be converted to Dask arrays for general Python workflows. BiocPy’s `DelayedArray` preserves the exact structure of Bioconductor’s delayed operations, allowing pipelines to be serialized and reconstructed across R and Python without eager evaluation or materializing a full copy of the matrix. Furthermore, `DelayedArray` integrates with the `tatami` C++ matrix interface, enabling downstream analysis packages to stream chunks directly from delayed or file-backed arrays during computation.

## 3 Cross-Language Data and Methods Interoperability

Providing compatible in-memory classes in Python is only part of cross-language integration. Workflows also require reliable ways to exchange serialized data on disk and guarantee that statistical algorithms behave consistently across languages. BiocPy addresses data exchange through two complementary mechanisms, direct reading and writing of R serialized (`.rds`) files via `rds2py` and schema-validated, language-agnostic persistence via `ArtifactDB`, and addresses algorithmic consistency through shared C++ libraries wrapped across R, Python, and WebAssembly.

### 3.1 Reading and Writing Serialized RDS Files

Many public genomic datasets and existing R pipelines rely on `.rds` files created via `saveRDS()`, which cannot be read or written natively in Python. To support direct `.rds` exchange without requiring an R installation, BiocPy includes `rds2py`[32], a library for reading and writing R’s binary serialization format. When reading `.rds` files, `rds2py` parses the low-level R object representation from disk into a lightweight dictionary structure and dispatches registered constructors to instantiate the corresponding BiocPy class, parsing atomic vectors and sparse matrix buffers directly into NumPy arrays and SciPy sparse matrices without requiring an intermediate R session. Conversely, `rds2py` can serialize Python data structures and BiocPy containers back into standard `.rds` files that can be loaded directly in R via `readRDS()`, allowing Python tools to integrate into existing R/Bioconductor workflows without changing file formats (see the executable code snippet).

### 3.2 Language-Agnostic Formats with ArtifactDB

While `rds2py` provides convenient `.rds` exchange with existing R scripts, R’s binary serialization format requires reading or writing entire objects in memory and is tied to R’s internal object representation rather than open storage standards. For language-agnostic archival storage and out-of-core access to large matrices, BiocPy integrates with the schemas and specifications defined by ArtifactDB[33], implemented by the `alabaster` suite in R[34] and the `dolomite` suite in Python[35].

Under this framework, complex Bioconductor and BiocPy objects are decomposed on disk into standard, language-neutral file formats: HDF5 for dense or sparse matrices and tabular data, and JSON for object metadata. An R user can stage a `SingleCellExperiment` or `MultiAssayExperiment` to a directory using `alabaster`, and a Python user can read that directory using `dolomite` to reconstruct the corresponding BiocPy object (and vice versa). When objects are loaded in either R or Python, assay matrices are wrapped as `DelayedArray` objects rather than read eagerly into memory, enabling out-of-core access to large datasets. This decouples on-disk data persistence from any single programming language. Consistency of the file representations is enforced by a validation library written in C++, ensuring that objects saved from either R or Python can be read faithfully in the other language (see the R and Python snippets).

### 3.3 Shared C++ Libraries as a Paradigm for Cross-Platform Method Development

Even when two languages share identical data structures, independently reimplementing statistical methods in R and Python frequently introduces subtle algorithmic discrepancies[36]. Maintaining separate codebases also doubles the engineering effort required to fix bugs, optimize performance, and validate edge cases. To address this, computational packages in the BiocPy ecosystem follow a shared C++ core architecture (Figure 2), which we recommend as a general development paradigm for bioinformatics method developers seeking to make their tools accessible across multiple languages.

**Figure 2:**
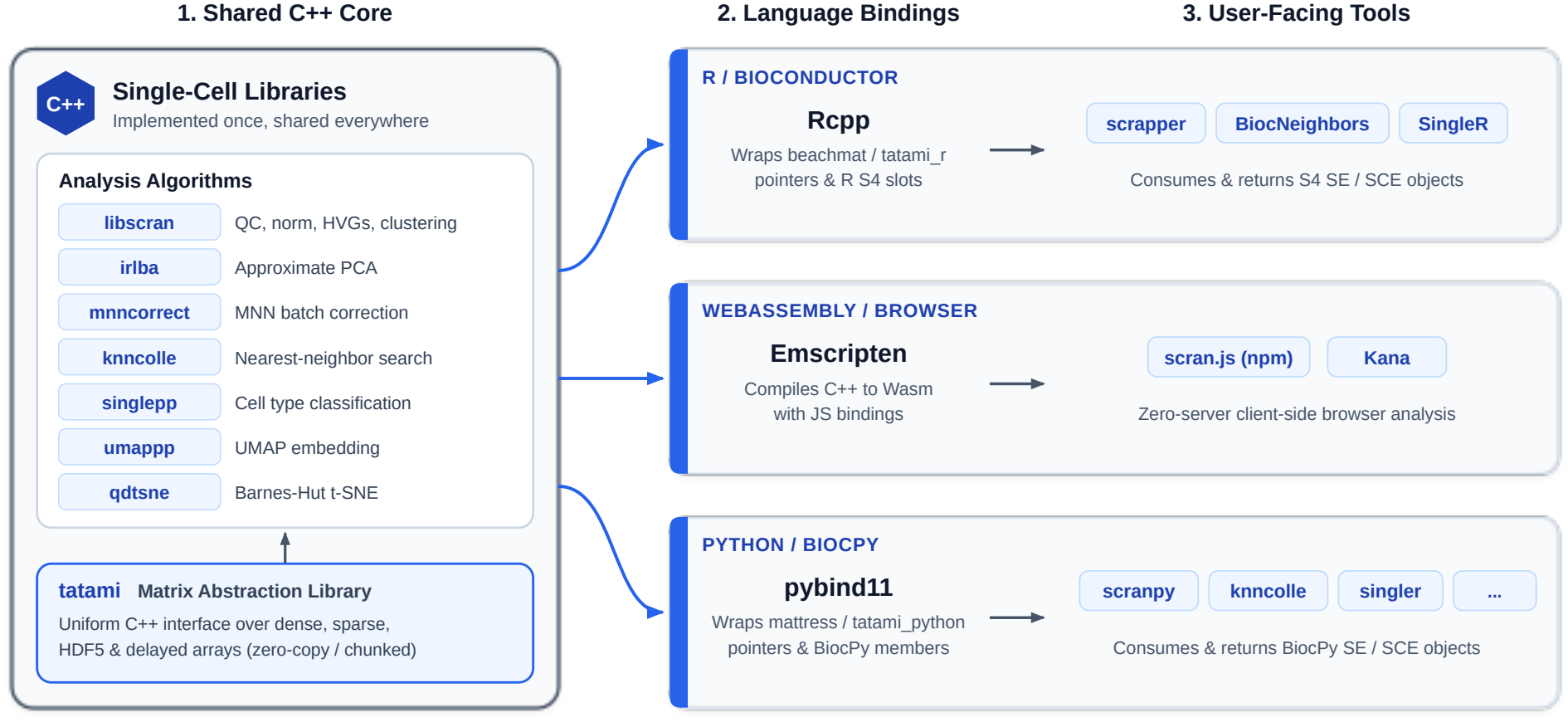
Shared C++ libraries enabling single-cell analyses across computational environments. Core C++ algorithms (`libscran`, `irlba`, `mnncorrect`, `knncolle`, `singlepp`, `umappp`, and `qdtsne`) build on the `tatami` matrix abstraction layer. These are wrapped via `Rcpp` (with `beachmat`/`tatami_r`) for Bioconductor, Emscripten[45] for WebAssembly (`scran.js`, Kana), and `pybind11` (with `mattress`/`tatami_python`) for BiocPy. This architecture ensures that high-level calls in R, Python, or JavaScript execute the same underlying C++ implementation, preventing algorithmic differences.

Under this paradigm, core algorithms are implemented once in C++ libraries. Matrix access is abstracted through the C++ `tatami` library[37], which provides a uniform iteration interface over in-memory dense and sparse arrays (accessed in place without copying) as well as delayed or HDF5-backed matrices (streamed in chunks without materializing the full matrix). High-level analysis libraries such as `libscran`[38], `knncolle`, and `singlepp`[39, 40] operate directly on `tatami` handles. Language-specific packages then wrap this shared core: R/Bioconductor packages use `Rcpp` (e.g., `scrapper`[41]) to pass R matrices and S4 slots into `tatami` and return `SingleCellExperiment` or `SummarizedExperiment` objects; Python/BiocPy packages use `pybind11` (e.g., `scranpy`[42] and `singler`[43], `knncolle`, and `mattress`) to pass NumPy arrays and SciPy sparse matrices to the same routines and populate BiocPy container members; and compiling the same C++ sources to WebAssembly[44] via Emscripten[45] powers browser-based applications such as Kana[46].

By pairing a language-neutral C++ computational layer with consistent high-level container semantics (`SummarizedExperiment` and `SingleCellExperiment`) in R and Python, method developers can write and test an algorithm once while exposing idiomatic APIs across ecosystems.

## 4 Case Studies

Combining native Bioconductor-compatible containers with shared C++ backends and language-neutral persistence enables end-to-end genomic workflows in Python as well as hybrid pipelines spanning R and Python. Below, we present three workflows illustrating range-based genomic overlap queries, cross-language access to curated data repositories, and multi-modal single-cell analysis.

### 4.1 Genomic Interval Operations

To illustrate range-based genomic workflows in Python, we consider integrating Bioconductor reference annotations with external peak-calling datasets. In Bioconductor, transcript and gene models are commonly distributed as SQLite-backed `TxDb` packages through `AnnotationHub` and `ExperimentHub`[47]. The `txdb`[48] Python package queries these SQLite databases directly and constructs BiocPy `GenomicRanges` and `GenomicRangesList` objects representing genes, transcripts, exons, and coding sequences (complemented by the `ensembldb`, `orgdb`, and `experimenthub` packages for querying Ensembl annotations, organism-level gene mappings, and hub resources). This allows Python users to access curated Bioconductor genome annotations directly without manually downloading and parsing raw GTF or GFF files.

Once the human reference (e.g., hg38) transcript coordinates are loaded as a `GenomicRanges` object, interval methods can be chained to define functional regions and query experimental intervals. For example, promoter regions around transcription start sites are computed, and ChIP-seq peak intervals are fetched from BEDbase[49] as a `GenomicRanges` object. Calling overlap and proximity methods (such as `find_overlaps()` or `subset_by_overlaps()`) identifies peaks overlapping promoter regions in a few lines of Python code while preserving sequence metadata and strand semantics (see the ChIP-seq interval snippet).

### 4.2 Cross-Language Data Repositories with scrnaseq and celldex

Shared data repositories often duplicate effort when datasets must be packaged separately for R and Python users. The `scrnaseq`[50, 51] package (providing curated public single-cell datasets) and the `celldex`[52, 53] package (providing reference expression profiles for cell type annotation) are both available in R and Python and store their underlying datasets in the ArtifactDB format. Because ArtifactDB decomposes each dataset into language-neutral HDF5 matrices and JSON metadata validated against shared schemas, a single versioned deposition can be served to multiple language ecosystems without maintaining separate file formats or conversion pipelines.

In R, `scrnaseq` and `celldex` reconstruct these artifacts via `alabaster` into Bioconductor `SingleCellExperiment` and `SummarizedExperiment` objects; in Python, the corresponding BiocPy packages use `dolomite` to load the exact same versioned artifacts into native BiocPy containers. Other programming languages (such as JavaScript or C++) can likewise access the same repositories through ArtifactDB-compatible readers, ensuring consistent dataset versions, feature annotations, and sample metadata across platforms.

### 4.3 Multi-modal Single-Cell Analysis using scranpy

To demonstrate how BiocPy connects these shared data repositories with cross-language statistical methods, we consider a single-cell analysis and annotation pipeline in Python. First, a curated single-cell dataset is fetched from `scrnaseq` as a `SingleCellExperiment` object. The dataset is then processed using `scranpy`[42], which performs quality control filtering, scaling normalization, variance modeling, principal component analysis, clustering, and multi-modal integration across primary RNA counts and auxiliary modalities (such as ADT counts in `alt_exps`). This matches the Bioconductor workflow implemented by R packages (such as `scrapper`) compiled with the same underlying C++ libraries[41, 54, 23].

Following preprocessing, cell types are annotated using the `singler` Python package[43] against a `celldex` reference profile, again sharing its C++ implementation with the R/Bioconductor `SingleR` package. All computed embeddings (PCA, t-SNE, UMAP), cluster assignments, and cell type labels are stored directly in the `reduced_dims` and `column_data` members of the `SingleCellExperiment` object for visualization with standard Python plotting libraries such as `seaborn` or `matplotlib` (complete Python and R workflow snippets are available in the paper repository).

## 5 Discussion

BiocPy provides a practical bridge between the R/Bioconductor and Python ecosystems, though several scope boundaries and trade-offs warrant consideration. First, BiocPy focuses on core data structures, serialization utilities, and analysis workflows (such as single-cell and interval operations); it does not aim to port the more than 2,000 specialized downstream statistical packages in Bioconductor. When an analysis requires specialized R-only modeling packages, BiocPy’s serialization tools (`rds2py` and `dolomite`/`alabaster`) allow data to be exchanged cleanly between R and Python steps in a multi-language pipeline. Second, BiocPy prioritizes strict dimensional validation, explicit member structures, and copy-on-write (`in_place=False`) semantics to mirror S4 behavior. While these guarantees help prevent silent data corruption in complex pipelines, they can require adjustment for Python users accustomed to unconstrained in-place mutation on dictionary- or DataFrame-backed objects. Finally, while `rds2py` supports reading and writing core Bioconductor classes out of the box, exchanging bespoke, package-specific S4 classes via `.rds` files requires registering custom parser and serializer mappings.

Beyond data exchange, the combination of BiocPy containers and shared C++ libraries (`tatami`, `libscran`) provides a practical template for method developers building cross-language bioinformatics tools. Rather than writing a method in R and later porting it to Python or vice-versa, developers can implement core algorithms in lower-level languages such as C, C++, or Rust against abstract matrix interfaces and expose thin wrappers that consume and return `SummarizedExperiment` or `SingleCellExperiment` objects in each language. Because the high-level container layout is preserved across Bioconductor and BiocPy, wrapper code remains minimal, documentation and tutorials translate directly across languages, and users in both ecosystems have consistent algorithmic implementations.

Looking ahead, formalizing a language-agnostic specification for Bioconductor’s core classes (analogous to W3C web standards or the Arrow columnar specification) would further strengthen cross-language interoperability. Such a standard would define the required members, structural invariants, coordinate conventions, and subsetting semantics for containers such as `GenomicRanges` and `SummarizedExperiment`, paired with a shared conformance test suite so independent implementations in R, Python, Julia, Rust, or WebAssembly can be verified against a common reference.

All packages in the BiocPy ecosystem follow consistent software engineering practices, including test suites, continuous integration, and standardized project scaffolding, to ensure long-term maintainability. By providing native Python implementations of Bioconductor’s core data structures alongside shared C++ computational backends and language-neutral storage formats, BiocPy reduces the friction of working across R and Python. Analysts and tool developers can represent genomic assays, intervals, and multi-omics datasets using a consistent conceptual model while taking full advantage of Python’s machine learning and data science libraries.

## 6 Data and Code Availability

The BiocPy ecosystem is open-source and released under the MIT license. All packages are published to the Python Package Index (PyPI). Source code, issue tracking, and documentation are maintained under the BiocPy GitHub organization (https://github.com/BiocPy), and contributor guidelines are available in the developer guide (https://github.com/biocpy/developer_guide).

## 7 Acknowledgements

We acknowledge Jean-Philippe Fortin, Vincent Carey, Hector Corrada Bravo, Michael Lawrence, and Mike Love for their support of the BiocPy effort. We thank Kevin Yang for his contributions to improving BiocPy, and Xiaosai Yao for reviewing and providing valuable feedback on this manuscript.

